# The *vgll3* locus controls age at maturity in wild and domesticated Atlantic salmon (*Salmo salar* L.) males

**DOI:** 10.1101/024927

**Authors:** Fernando Ayllon, Erik Kjærner-Semb, Tomasz Furmanek, Vidar Wennevik, Monica F Solberg, Geir Dahle, Geir Lasse Taranger, Kevin A Glover, Markus Sällman Almén, Carl J Rubin, Rolf B Edvardsen, Anna Wargelius

## Abstract

Wild and domesticated Atlantic salmon males display large variation for sea age at sexual maturation, which varies between 1-5 years. Previous studies have uncovered a genetic predisposition for age at maturity with moderate heritability, thus suggesting a polygenic or complex nature of this trait. The aim of this study was to identify associated genetic loci, genes and ultimately specific sequence variants conferring sea age at maturity in salmon. We performed a GWAS using a pool sequencing approach (20 individuals per river and trait) of salmon returning to rivers as sexually mature either after one sea winter (2009) or three sea winters (2011) in six rivers in Norway. The study revealed one major selective sweep, which covered 76 significant SNP in which 74 were found in a 370 kb region of chromosome 25. Genotyping other smolt year classes of wild salmon and domesticated salmon confirmed this finding. Genotyping domesticated fish narrowed the haplotype region to four SNPs covering 2386 bp, containing the *vgll3* gene, including two missense mutations explaining 33-36% phenotypic variation. This study demonstrates a single locus playing a highly significant role in governing sea age at maturation in this species. The SNPs identified may be both used as markers to guide breeding for late maturity in salmon aquaculture and in monitoring programs of wild salmon. Interestingly, a SNP in proximity of the *VGLL3* gene in human (*Homo sapiens*), has previously been linked to age at puberty suggesting a conserved mechanism for timing of puberty in vertebrates.

**Author summary:** For most species the factors that contribute to the genetic predisposition for age at maturity are currently unknown. In salmon aquaculture early maturation is negative for the growth, disease resistance and flesh quality. In addition, using populations of salmon selected to mature late may limit the genetic impact of aquaculture escapees, as these late maturing fish are more likely to die before they reach maturity. The aim of this study was to elucidate the genetic predisposition for salmon maturation. We determined the sequences of genomes from Atlantic salmon maturing early and late in six Norwegian rivers. This methodology enabled us to identify a short genomic region involved in determining the age at maturity in Atlantic salmon. This region has also previously been linked to time of puberty in humans – supporting a general mechanism behind age at maturity in vertebrates. The results of this study may be used to breed salmon that are genetically predisposed to mature late which will improve welfare and production in aquaculture industry and aid in the management of escaped farmed salmon.

## Introduction

Both wild and domesticated farmed populations of Atlantic salmon (*Salmo salar* L.) show large phenotypic variation for sea age at sexual maturity [1]. Salmon males can stay in the sea from 1-5 years before they initiate sexual maturation and return to their native river to spawn, while females usually return to the river after 2-3 years in the sea. This variation in age at sexual maturity, especially for males, can also be observed in salmon aquaculture, where precocious puberty of males has been a major problem due to negative effects on somatic growth, flesh quality, animal welfare and susceptibility to disease [1]. Early maturation in farmed salmon can also increase the risk of further genetic introgression of escaped salmon in wild populations [2,3], as maturing fish will have a higher likelihood of migrating to a nearby river to spawn. Immature fish on the other hand will more likely migrate to sea where mortality is high before reaching maturity [4]. The impact of precocious maturity in aquaculture farms has been greatly reduced using continuous light treatment during winter-time, which appears to override the genetic predisposition for early puberty and thus exemplifying the plastic nature of onset of maturation [1]. Further improvement of selective breeding for late puberty is, on the other hand, hampered as the current production conditions, using continuous light, can mask genetic traits related to early puberty, making it difficult to gain further improvements of this trait by classical breeding protocols. In recent years sea temperatures have been higher than usual in Norwegian waters [5], which might override the effect of continuous light treatment in delaying maturation, causing unwanted male precocious maturation at the postsmolt stage [6], and possibly also increase incidence of unwanted sexual maturation after one sea winter. Increased water temperature associated with climate change demands better breeding program, which ultimately leads to production fish with a more robust trait for late maturation.

Salmonids in general display moderately high heritability for age at sexual maturity [7-11] and QTLs relating to this trait have been identified [12]. Also three recent papers used single nucleotide polymorphism (SNP) arrays to identify regions under selection for sea age at puberty in both, an aquaculture strain using a low density SNP array [13,14] and in wild populations using a high density SNP array [15]. These three reports revealed association of the trait to multiple loci but gave no clear answer regarding possible mechanisms, genes and regions behind age at puberty. Previous studies screened the genome for loci under selection using a limited set of SNPs, which may exclude the causative variants [16]. The recent salmon genome assembly [17] gives the opportunity to map massive parallel sequencing reads to the assembly, thereby enabling genome-wide detection not only of novel SNPs, but also small indels and structural variation [18]. Hence, the use of sequencing allows prediction of genetic variants effect on regulatory regions and genes, which may link traits with new biological mechanisms and open venues for subsequent functional studies.

This study aimed to elucidate the genes and genomic regions that regulate sea age at puberty in male Atlantic salmon. To investigate this trait, we performed genome resequencing of scale samples from wild salmon from six rivers in Western Norway, returning as sexually mature to rivers either after one or three years at sea (Fig 1). Using this approach we identified a region on chromosome 25 (Chr 25) harboring a dense set of significant SNPs in a stretch of 370 kb. These results were also confirmed in other year classes and in domesticated salmon. To conclude this study successfully demonstrated the importance of one single genomic region in determining age at maturity in male salmon.

**Fig 1.**
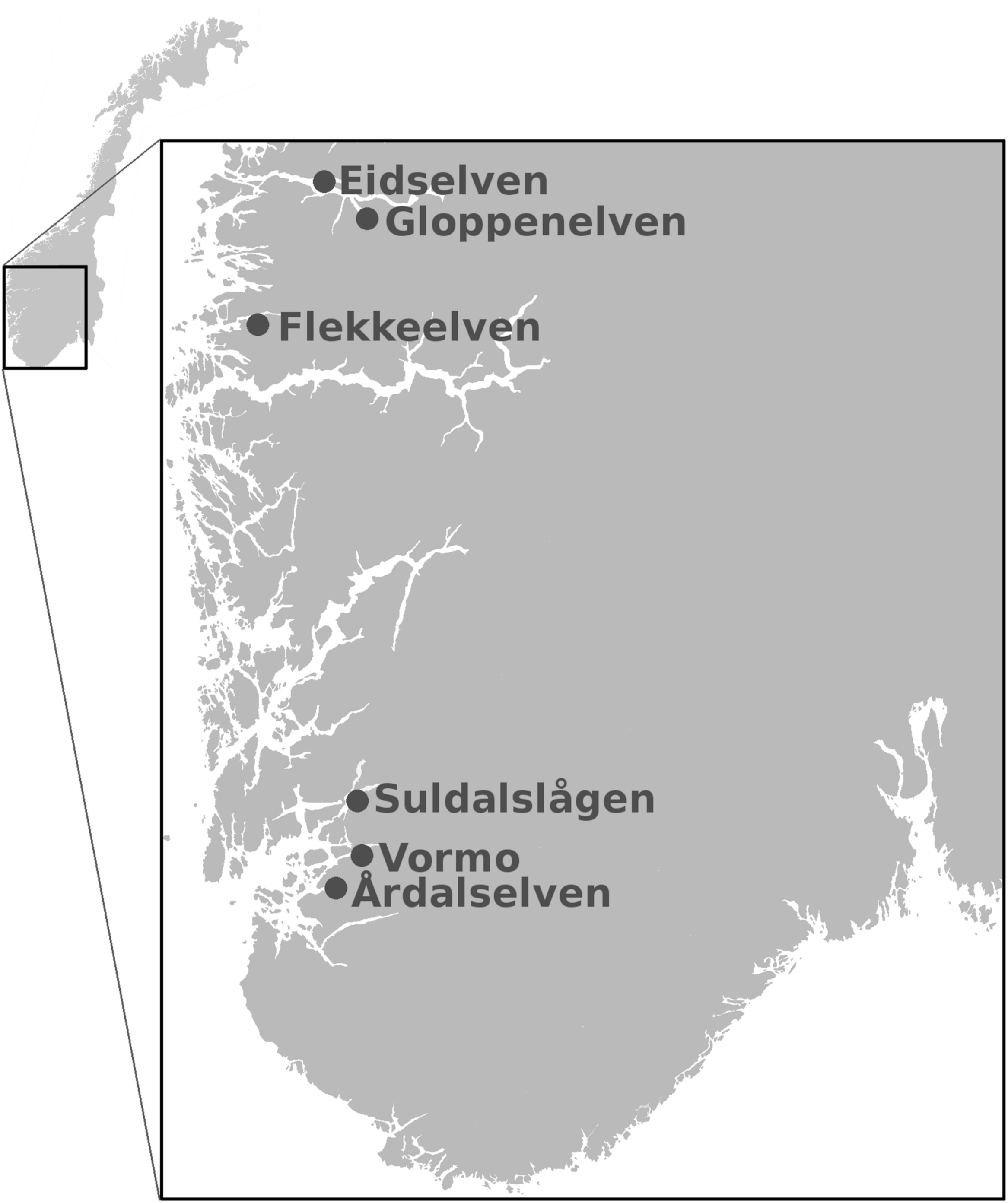
Geographical location of salmon rivers used. Map of Norway and a magnification of Western Norway showing rivers used in the experiment including three rivers in Sogn og Fjordane county; Eidselven, Gloppenelven and Flekkeelven, and three rivers in Rogaland county; Suldalslågen, Vormo and Årdalselven. Derivative of https://commons.wikimedia.org/wiki/File:Norway_municipalities_2012_blank.svg, licensed with CC-BY-SA-2.0.

## Results and Discussion

To find SNPs associated with age at maturation in salmon males, we sequenced 20 salmon per river and sea winter age (1SW and 3SW). This number of individuals in each pool has been shown to be sufficient to identify causative SNPs for a trait in *Drosophila melanogaster* [19]. Mapping our data yielded a 12.32X mean coverage (0.24 SE) of unique mapped reads per river and sea age at puberty (S1 Fig). This depth of coverage is similar to what has been used in other successful genome wide association studies (GWAS) by pool sequencing in vertebrates, including pig (*Sus scrofa*) and chicken (*Gallus gallus*) [20,21]. We have mapped the salmon sequences to the most recent salmon genome assembly (AKGD00000000.4). Within this assembly, 34% of the genome has not been assigned to chromosomes, probably due to a high number of repetitive sequences. This unassigned part of the genome harbored only 1% of our uniquely mapped reads (S1 Fig). SNP calling revealed altogether 4,326,591 SNPs in all sea ages and rivers.

Comparing 1SW and 3SW allele frequencies using the Cochran-Mantel-Haenszel (CMH) test revealed 155 SNPs significantly associated (0.1% FDR) with sea age at puberty (Fig 2A, S1 Table). Several single significant SNPs were found on chromosomes 1-7, 9-24 and 27-29, although these were not among the most significant SNPs (Fig 2A, S1 Table). In a previous QTL study for precocious parr maturation the trait was shown to be linked to Chr 12 [22]. Chr 12 has also been associated with sea age at maturation in another study [23]. In a GWAS, using a 6.5 kb SNP chip, the trait of 1SW maturation or “grilsing” was found to be weakly linked to both Chr 12 and Chr 25 [13]. In our study, 74 of the 155 (48%) significant SNPs were located in a region on Chr 25, covering ∼370kb (Fig 2B). From our data we conclude that in Western Norway a single selective sweep on Chr 25 has a large effect on sea age at maturity while other regions in the genome might contribute to a lesser degree. This is in contrast to earlier reports showing a more polygenic **nature** of this trait, with contributions from several genomic regions [13,14,23]. However, one previous model based study also suggested that age at maturation could be regulated by a stable genetic polymorphism, in accordance with our current findings [24].

**Fig 2.**
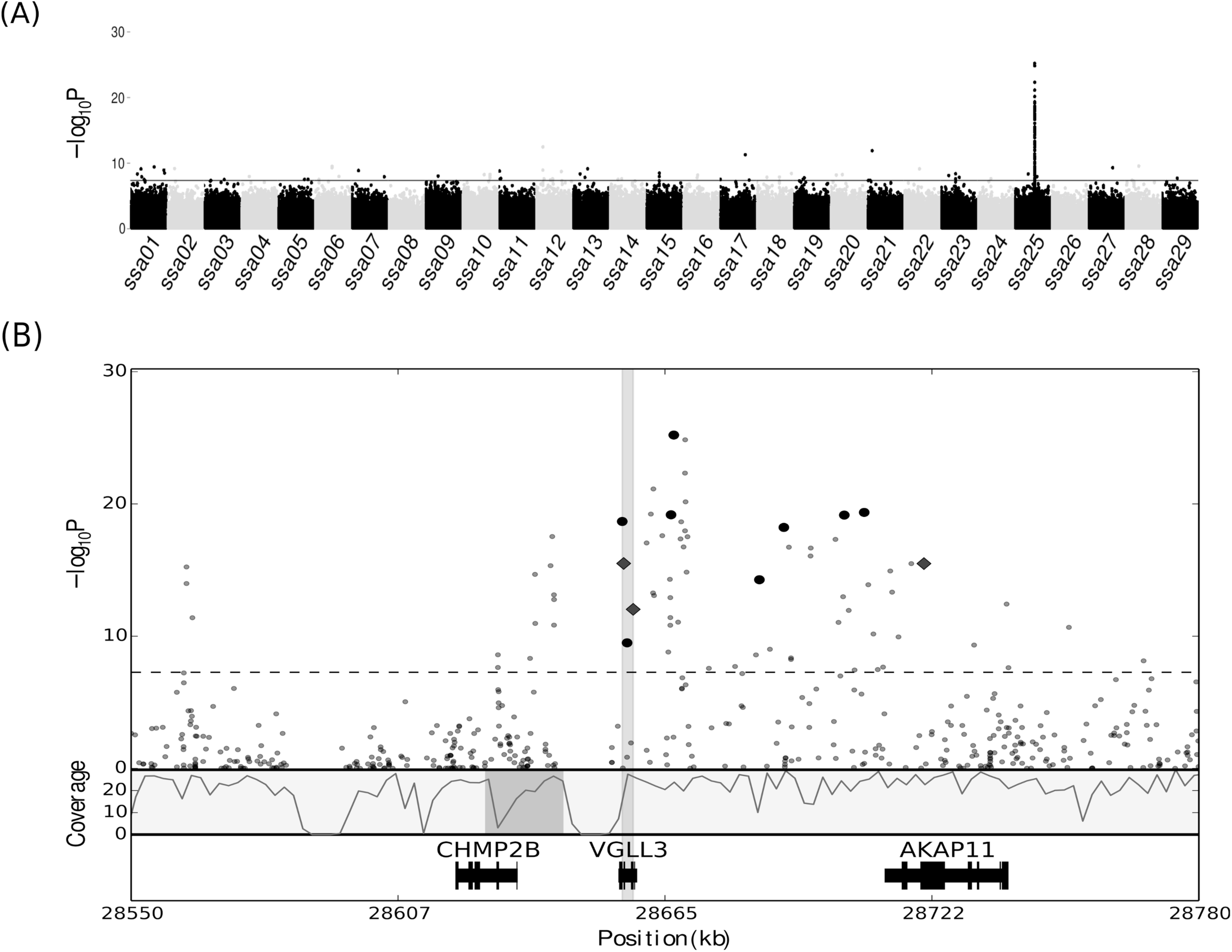
Identification of a selective region conferring age at maturity in Atlantic salmon. Manhattan plot of SNPs associated with age at maturity. The x-axis presents genomic coordinates along chromosomes 1-29 in Atlantic salmon. On the y-axis the negative logarithm of the SNPs associated p-value is displayed. All SNPs above the solid horizontal line in the plot are significantly (p < 5.2e^-8^, 0.1% FDR) associated with the trait. (B) Magnification of the 370kb region showing 230 kb on Chr 25 (28550-28780 kb) including 62 significant SNPs. The SNPs are represented by black dots, where the 11 large dots indicate SNPs used in genotyping assays. The square dots indicate three missense mutations in *vgll3* and *akap11*. On the upper y-axis the negative logarithm of the SNPs associated p-value is displayed. The dashed line within the plot indicates the significance threshold (0.1% FDR). Below the SNP illustration, the lower y-axis shows average read depth of coverage (using 2 kb windows). Genomic organization of the three genes found in the region is illustrated in the bottom track. The x-axis is showing the location of the region in the Chr 25 and covers 28550-28780 kb. The grey area around *vgll3* demarcates the shorter region discovered in the domesticated strain. The dark grey are in the coverage track is showing the misplaced contig in the most recent salmon genome release containing exon 1 and 2 of *chmp2b*.

To ascertain genotypes of single individuals for SNPs associated with sea age at maturity we designed a Sequenom assay for 11 of the most significant SNPs in the selective sweep found in Chr 25 (S2 Table). The genotyping results confirmed a strong association between allele frequencies and age at maturity (S2 Fig). To characterize haplotypes using the 11 assayed SNPs, we performed a pairwise disequilibrium analysis on all samples that had been sequenced [25]. This analysis revealed two dissimilar haplogroups comprising 11 haplotypes in one block (Fig 3A). One and five of these haplotypes showed significant association with maturing early and late, respectively. The significant 1SW haplotype explained 54% (β-value -1.0, p-value =3.88e-40) of the phenotypic variance for this trait. The most significant 3SW haplotype explained 21% (β-value 0.66, p-value=3.98e-13) of the variation in age at maturity, the other four 3 SW haplotypes explained the 1.9, 2.2, 3.6 and 3.7%, adding up to 32.4% of the variance of the age at maturity in 3SW haplotypes. The genotyping data clearly confirmed our findings from the pool re-sequencing study and further supported that this locus exerted a large effect on the trait.

**Fig 3.**
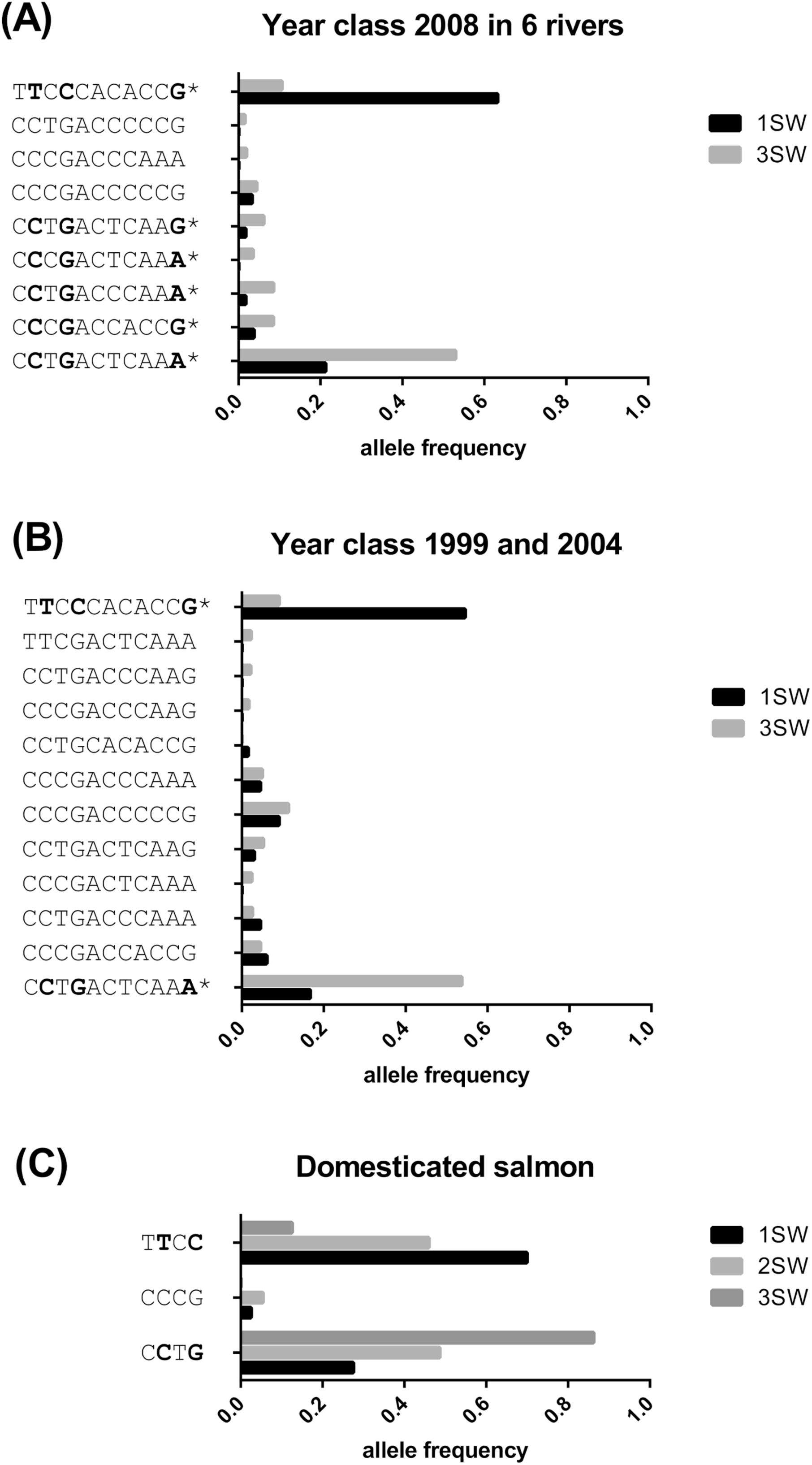
Haplotype frequencies in different year classes in wild and domesticated salmon. (A) Haplotype frequency associated with either 1SW (black bars) or 3SW (dark grey bars) in male Atlantic salmon for six rivers in Western Norway from year class 2008. (B) Haplotype frequencies associated with either 1SW (black bars) or 3SW (dark grey bars) male Atlantic salmon in year classes 1999 from Eidselva and 2004 from Suldalslågen. (C) Haplotype frequencies linked to time at maturity in the domesticated Mowi strain maturing after 1 (black bars), 2 (grey bars) or ≥ 3 (dark grey bars) years in sea water. In all graphs the x-axis indicates frequency of that trait for the identified haplotype, while the y-axis presents the haplotype block obtained from the genotype assay. * Indicates that the haplotype was significantly linked to the trait. The bold bases in the haplotypes are indicating missense mutations.

In samples from the pool sequencing we identified haplotypes associated with sea age at maturity in the 2008 year class (year of migration to sea). These fish have possibly been exposed to similar environmental conditions during their early stay in the sea, therefore showing a selection for those conditions as postulated by several previous studies in salmon [26-28]. To investigate this we identified genotypes using the SNP assays in other year classes: 1999 for Eidselva (20 1SW, 8 3SW) and 2004 for Suldalslågen (13 1SW, 13 3SW). Allele frequencies derived from the 11 SNP assays showed correlation to the 1SW and 3SW trait also in these year classes (S3 Fig). Haplotype association analysis of these year classes again revealed two significant haplotypes also found in the 2008 year class (Fig 3B). In the 1999 and 2004 year classes 44% (β-value -0.96, p-value=3.77e−08) and 22% (β-value 0.66, p-value=2.85e-04) of the phenotypic variation for age at maturity was explained by the 1SW or the 3SW haplotype, respectively. We thus conclude that genotypes at a single locus strongly influence sea age at maturation independent of year class across multiple salmon populations in Norway.

Sea age at maturity can be significantly altered in salmon by modulating both light and temperature [1,29,30]. As a consequence, current aquaculture production methods include the use of constant light during the winter months to inhibit or reduce the incidence of early sexual maturation. In turn, this means that selection for a genetic predisposition to mature late has been relaxed since the mid90s, when light regimes became part of the standard rearing procedure in Norwegian salmon farming, and partially concealing the genetic predisposition to mature early. We were also interested to see how much the identified genetic trait contributed to the sea age at maturity trait in domesticated farmed salmon males, since wild salmon live in a different environment including different feed availability and water temperature that may trigger time of male puberty differently. To assay the linkage between phenotype and genotype for sea age at maturity in a domesticated strain, we utilized DNA from sexually maturing salmon from four different families of the Mowi strain. This strain has been in aquaculture for at least ten generations and has been selected for a variety of traits including growth and late maturation [31-33]. Mowi was originally obtained from a range of large wild salmon populations from Western Norway in 1969, and has later been bred using a four-year life cycle. The long life cycle breeding has thereby probably increased the allele frequency for the late maturity trait. In this common garden experiment with the Mowi strain, fish were grown under natural light conditions in sea cages where males were maturing after 1, 2 or 3 or more years in sea. Haplotype analysis of these fish (n=97) revealed a shorter haplotype, consisting only of four SNPs, covering only 2386 bp in the 5’ end of the region assayed (Fig 3C, S4 Fig). These data clearly demonstrate that age at puberty can be explained by SNPs in this region also in a domesticated strain in culture for more than ten generations.

Altogether the experiments clearly show that the selective sweep on Chr 25 significantly contributes to sea age at maturity both in wild and domesticated salmon. Gene prediction in this area revealed three genes; *charged multivesicular protein 2B* (*chmp2B*), *vestigial-like protein 3* (*vgll3*) and *a-kinase anchor protein 11* (*akap11*, Fig 2B). From the analysis of domesticated salmon we could decrease the area of selection to a 2.4 kb region covering only *vgll3*. From our genotyping assay on domesticated samples we could with certainty reduce the region under selection in the downstream region of the *vgll3* locus since we had genotyped several SNPs in this area. We can however not exclude that the upstream region of the *vgll3* locus contained SNPs contributing to the haplotype since this area was not represented in our genotyping assay due to a large gap in the genome at this region (Figure 2B). The 2.4 kb region contained two missense mutations in *vgll3;* at amino acid (aa) 54 and 323. The age at maturity trait was strongly associated to the genotype for these mutations showing 36% (nt. 28656101 Chr25, β-value -0.61, p-value=9.80e-07) and 33% (nt. 28658151 Chr 25, β-value -0.60, p-value=3.77e-08). The haplotypes associated with the 3SW trait encode a Thr and a Lys at these positions whereas the haplotype associated with 1SW encodes a Met and an Asp. Our analysis could not conclude whether these missense mutations are causative for the sea age at maturity trait, but since they occurred consistently together in the material we cannot rule out whether both or other non-coding variants at this locus are involved in age at maturity phenotype. It is also known from other studies that co-occurring amino acid changes can confer a phenotype [34]. The Vgll3 protein functions as a cofactor for the TEAD family of transcription factors [35]. The transcription factor binding region spanning aa105-aa134 in Vgll3 does not include any of the aa changes discovered which suggests that any direct binding differences between 1SW or 3SW fish are unlikely. It is thus difficult to predict how these amino acid changes affect the protein. At this point we cannot elucidate whether it is these missense mutations or other SNPs outside coding regions, which confer the trait. The question about the ancestral and derived alleles remains elusive, but we surveyed sequences from other salmonids for information about the amino acid variants and found that both brown trout (*Salmo trutta* L.), rainbow trout (*Oncorhynchus mykiss*) and arctic char (*Salvelinus alpinus*) all harbor the 3SW variants of the amino acids. In addition, we ran an allelic discrimination assay on five individuals from the Swedish landlocked Atlantic salmon population, Gullspång (landlocked for 10,000 years), all carrying only the 3SW (Thr-Lys) amino acid variant. This indicates that the 3SW version of the Vgll3 is ancestral and that the 1SW (Met-Asp) is derived.

In humans the VGLL3 locus has been linked to age at maturity or puberty by a SNP in close proximity of the gene [36], strengthening our notion that the salmon Vgll3 protein is involved in age at puberty in fish. Regarding the function of this protein in controlling age at maturity, it is known that Vgll3 is involved in the inhibition of adipocyte differentiation in mouse (*Mus musculus*) [37]. Changes in fat metabolism may be partially causative for changes in age at maturity, since increased adiposity has previously been linked to maturation in salmon [38-41]. In studies in rodent testis, *vgll3* transcripts have been associated with differential expression during the early stages of steroidogenesis in the embryonic testis [42], suggesting a role in testis maturation. Further functional studies of this protein and adjacent regulatory regions will confirm if the previous study in humans and our study have actually revealed a universal regulator of age at maturity in vertebrates.

The most significant SNPs were located in the *vgll3* locus but two neighboring genes, *chmp2B* and *akap11*, also contain several significant SNPs (Fig 2B). One of these, a missense mutation in *akap11* translates to a Val in 1SW and a Met in 3SW at aa 214. AKAP11 is involved in compartmentalization of cyclic AMP-dependent protein kinase (PKA). This aa AKAP11 is not located in any of the known functional domains related to PKA [43]. AKAP 11 is highly expressed in elongating spermatocytes and mature sperm in human testis and is believed to contribute to cell cycle control in both germ cells and somatic cells. There are no reports clearly linking this protein to age at maturity but future functional studies will reveal if that is the case. *Chmp2B* did not contain any missense mutations but upon manual review of this region we detected a misplaced 16,885 bp region in the Chr 25 containing exon 1 and 2 of *chmp2B*. This region also carried many significant SNPs which were probably associated with the selective sweep. When this region was placed in proximity of the gene (dark grey box in Fig 2B) it became clear that many significant SNPs were near the *chmp2b* gene. This gene encodes a protein belonging to a protein complex which is involved in protein endocytosis [44] (Fig 2B). In humans CHMP2B is known to be essential for the survival of nerve cells and is linked to both dementia and ALS [45-47]. It is well known that the neural system works as a gatekeeper in controlling age of puberty, also in fish [48] but whether Chmp2B is involved in the regulation of puberty remains to be elucidated.

In this study we performed a GWAS by genome re-sequencing with the aim to screen the genome of Atlantic salmon for loci regulating age at maturity in males. By investigating late and early maturing male fish from six rivers in Western Norway we demonstrated that the sea age at maturity trait was strongly associated with sequence variation at one locus on Chr 25. The haplotype associated with late maturity can be used for selective breeding on individuals predisposed for this trait, thereby possibly reducing the incidence of negative phenotypes associated with early maturation of males in salmon aquaculture. This study also shows that certain haplotypes significantly contribute to the sea age at puberty, and may therefore be implemented as markers in the management of wild salmon populations in the face of changing environmental conditions such as increased sea temperatures. Significantly, this study and a previous study in humans [36], suggests a conserved role of the *Vgll3* protein in timing of puberty in vertebrates.

## Materials and Methods

### Samples and sampling

The samples of wild salmon upon which this study is based were collected by Rådgivende Biologer AS, Bergen, Norway (http://www.radgivende-biologer.no). Scales were taken from dead salmon fish that had been captured by anglers during the fishing season. In this manner, samples of wild salmon were acquired from six rivers in Western Norway; Eidselva, Gloppenelven, Flekkeelven, Årdalselva, Suldalslågen and Vormo (Fig 1). In order to minimize the potential influence of environmental variation on the sea age at maturity we used fish from the same smolt year class sampled as 1SW fish (returning to river 2009) and 3SW fish (returning to river 2011). Each river was represented by 20 1SW and 20 3SW males. For the genotyping assay we also included two other year classes from Eidselva and Suldalslågen. From Eidselva we retrieved scales from 20 1SW males from year 2000 and 8 3SW males from 2002. From Suldalslågen we obtained scales from 14 1SW and 14 3SW from years 2005 and 2007, respectively.

In addition to samples of wild salmon, we investigated age at maturity in four full sibling families of domesticated salmon from the Norwegian Mowi strain maturing at 1SW, 2SW or older. These fish were obtained from an ongoing study at the Matre Aquaculture Research station where they were reared in a common garden design in sea cages without the use of continuous light, i.e. under ambient light only. Before transfer to sea cages, fish were sedated (0.07 gL^-1^, Finquel®, ScanAqua), adipose fin clipped and PIT (passive integrated transponder) tagged. Fin clips, preserved on 95% ethanol, from a total of 97 fish maturing at different sea ages were included in this study. The four families consisted of 36, 24, 13 and 24 fish per family. The parents were also included in the study.

### DNA extraction and PCR-based sdY test

Total DNA from selected individuals was purified from 2 to 3 scales using Qiagen DNeasy Blood & Tissue Kit (Qiagen, Hilden, Germany) according to the manufacturer’s recommendations. Sex of all samples used herein was validated by a PCR-based methodology aimed to detect the presence of the *sdY* gene [49,50]. Individuals showing amplicons of exon 2 and 4 were designated as males. As a positive PCR control and for species determination we used the presence of the 5S rRNA gene [51]. PCR amplifications were performed using reaction mixtures containing approximately 50 ng of extracted Atlantic salmon DNA, 10 nM Tris–HCl pH 8.8, 1.5 mM MgCl_2_,50 mM KCl, 0.1% Triton X-100, 0.35 μM of each primers, 0.5 Units of DNA Taq Polymerase (Promega, Madison, WI, USA) and 250 μM of each dNTP in a final volume of 20 μL. PCR products were visualized in 3% agarose gels.

### Library preparation, sequencing and mapping

Following fluorometric quantification, equal amounts of DNA from ten males were pooled to generate paired-end libraries using the Genomic DNA Sample Preparation Kit (Illumina, CA, USA) according to manufacturer’s instructions. Libraries were sequenced on the Illumina HiSeq2000 platform (Illumina, CA, USA) at the Norwegian Sequencing center (http://www.sequencing.uio.no/, Oslo, Norway). In each sequencing lane we used pools of 10 fish from each sea age and river which made a total of 24 lanes sequenced in the whole experiment (6 rivers, 2 replicates per sea age). Raw sequence data has been deposited at SRA with BioProject Accession number PRJNA293012. Library quality control was conducted to ensure that all the samples fulfilled the quality standards (FastQC -http://www.bioinformatics.babraham.ac.uk/projects/fastqc/). Adapter and quality trimming of FastQ format reads were carried out using Cutadapt [52]. All 24 libraries containing on average 361821757 (± 4956053) paired end reads were approved for further analysis and aligned to the most recent salmon genome release (Acc. No. AGKD0000000.4) using Bowtie2 (v.2.1.0) [53]. Entire read alignment with no soft clipping was required by setting Bowtie 2 to the end-to-end mode. Seed length during alignment was set to 18, allowing only 1 mismatch. Interval function between seed substring during multiseed alignment was defined by the following variables: S,1,1,1.5. Maximum number of ambiguous characters was set by the following function parameters: L,0,0.1. Minimum alignment score was governed by the function L,-0.6,-0.4. Only unambiguously mapped reads (mapping quality score greater than 20) were retained for downstream analysis.

### SNP calling, annotation and statistical analysis

To improve the sensitivity to detect rare alleles, biological replicates in the dataset were bioinformatically fused using SAMtools merge, producing a single BAM file for each river and maturation stage [54]. SNPs were called using the Mpileup command in SAMtools. The resulting file was then recoded for use in the PoPoolation2 pipeline (v.1.2.2) [55]. A Minimum base quality threshold of 20 was established in order to remove ambiguously mapped reads and low quality bases. The Cochran-Mantel-Haenszel test for repeated tests of independence for every SNP was performed using the PoPoolation 2 package (cmh-test.pl and R based custom script) in order to detect significant differences (0.1% FDR) in allele frequencies between 1SW and 3SW pools [56]. For each merged sample of 20 fish, the parameters min-count was set to a value of 10 whereas the min-coverage and max-coverage were set to 7 and 42, respectively (5-95% percentile). To annotate the salmon genome (AKGD00000000.4), Augustus gene prediction software was trained using PASA gene candidates by mapping salmon ESTs from NCBI to the salmon genome assembly with PASA [57,58]. The Augustus de novo gene prediction contained coding sequences without UTRs. The genes were validated by RNASeq from both Atlantic salmon [59] and rainbow trout [60] and annotated with Swissprot. Significant SNPs were functionally annotated to predict variant effect (custom R and Python scripts). Bioinformatical analysis identified a 16,885 bp region in Chr 25 (position 28907421-28924305) which contained the first two exons of the gene *chmp2b* in addition to several significant SNPs. In a previous version of the genome assembly (AKGD00000000.3) this region existed as a single contig, and has presumably been inserted into the wrong chromosomal region in the most recent genome assembly. We corrected this by reverse-complementing the region and inserting it in the gap between the third exon of *chmp2b* and *vgll3* in Chr 25 (position 28626249-28643134) placing the exons of *chmp2b* in coherent order. Genotype/phenotype association analysis was performed using Plink v1.8 [25]. Selected SNPs were tested for association using asymptotic significance values (likelihood ratio test and Wald test 1%FDR [56]). Whenever possible, asymptotic haplotype-specific association tests were performed in order to establish the percentage of the phenotypic variation that could be explained by the detected haplotypes.

### Sequenom assays

Eleven of the most significant SNPs (S1 Table) identified in the putative selective sweep (Fig 2B) were used to design a Sequenom assay to genotype 296 Atlantic salmon individuals belonging to the six natural populations sampled (year classes 1999, 2004 and 2008) and 97 fish belonging to the 4 families from the domesticated strain. Genotyping was conducted on a Sequenom MassARRAY analyser (San Diego, CA, USA). A complete list of Primers and extension primers used are found in the S2 Table.

### Ethics Statement

Scale samples from wild salmon were collected by local anglers during the fishing season, thus no permits/licenses regarding the collection of these samples were required by the research team. Samples from domesticated fish were retrieved from an ongoing study at Matre Aquaculture Research station (IMR), where the experimental protocol (permit number 4268) had been approved by the Norwegian Animal Research Authority (NARA). Welfare and use of these experimental animals was performed in strict accordance with the Norwegian Animal Welfare Act of 19^th^ of June 2009, in force from 1^st^ of January 2010.

## Acknowledgements

We would like to acknowledge the expert advice of Eva Andersson, Rüdiger W Schulz, Jan Bogerd, Tom Hansen, Ketil Malde and Per Gunnar Fjelldal during the project. We are also grateful to Anne-Grethe Sørvik, Stig Mæhle, Sara Olausson, Sven Leininger, Amèlie Juanchich, and Lene Kleppe for providing laboratory support. Special thanks to Ivar Helge Matre and Lise Dyrhovden for providing excellent assistance in taking care of the salmon experiment performed at Matre aquaculture station. We would like to thank Rådgivende Biologer Bergen A/S for selling us the wild salmon samples.

## Supporting information

**S1 Fig. Mapping statistics**

(A) Average read depth in all pooled samples (x-axis). The y-axis is showing average percent of the genome covered, with error bars. (B) Average number of uniquely mapped sequences (y-axis) on each chromosome (x-axis), with error bars. “Scaffolds” refers to unplaced contigs in the current genome assembly.

**S2 Fig. Allele frequencies in year class 2008**

Frequencies of the late maturation allele in year class 2008 are shown for 1SW (green bars) and 3SW (blue bars) fish. The position of each SNP in Chr 25 is shown in the leftmost part. Above each bar the number genotyped fish is indicated. The y-axis shows the allele frequency between 0 and 1. Abbreviations; Ard - Årdalselven, Eid - Eidselven, Fle - Flekkeelven, Glo - Gloppenelven, Sul - Suldalslågen and Vor - Vormo.

**S3 Fig. Allele frequencies in year class 1999 and 2004**

Frequencies of the late maturation allele in Eidselva year class 1999 and Suldalslågen yearclass 2004 are shown for 1SW (green bars) and 3SW (blue bars) fish. The position of each SNP in Chr 25 is shown in the leftmost part. Above each bar the number genotyped fish is indicated. The y-axis shows the allele frequency between 0 and 1. Abbreviations; Eid - Eidselven, and Sul - Suldalslågen.

**S4 Fig. Allele frequencies in domesticated salmon**

Frequencies of the late maturation allele in farmed salmon (Mowi strain) strains, maturing either after 1SW (green bars), 2SW (yellow bars) or after 3SW or more in sea water (blue bars). The position of each SNP in Chr 25 is shown in the leftmost part. Above each bar the number genotyped fish is indicated. The y-axis shows the allele frequency between 0 and 1.

**S1 Table. Significant SNPs**

Table of SNPs significantly associated to age at maturity. The table includes scaffold names, chromosome position, reference/alternative (Ref/Alt) Alleles, Average Coverage (+/-SE) and P-values.

**S2 Table. Primers and extension primers used in the Sequenom assay**

## Data reporting

Sequenced pool material submitted to SRA (Bioproject number PRJNA293012)

